# Protocol for the Gene Dictionary Tool (GDT) to create and implement gene dictionaries across annotated genomes

**DOI:** 10.1101/2025.06.15.659783

**Authors:** Breno Dupin, Matheus Sanita Lima, Alexandre Rossi Paschoal, David Roy Smith

## Abstract

The Gene Dictionary Tool is a protocol for the creation and implementation of gene dictionaries across annotated genomes. Through a Python CLI (Command Line Interface), the user identifies what annotations are missing in the inputted dictionary. Via two Jupyter Notebooks, the user builds a gene dictionary based on attributes retrieved from inputted GFF3 files. The final output, a .gdict file, is Findable, Accessible, Interoperable, and Reusable (FAIR). We showcase GDT’s functionalities by creating a gene dictionary for fungal mitochondrial genomes.

## Before you begin

The deluge of publicly available Next Generation Sequencing (NGS) transcriptomic data has been dubbed a “goldmine for organelle research”^1^, and a similar conclusion can be drawn from DNA-Seq experiments^2^. Given this wealth of data, organelle genomes currently are among the most sequenced type of chromosomes^3^. Although there exist biases in taxonomic sampling^4^, comparative analyses of organelle genomes have informed all-encompassing theories of genome evolution^5^, shed light on what genome features explain organelle gene retention^6^, and shown how selection can act on some of these cytoplasmic genetic systems^7^. Even with such popularity and scientific importance, the field of organelle genomics suffers from a lack of gene nomenclature standardization^8^. Attempts to bring order and consistency in organelle gene nomenclature date back to at least 1983^9^ and, currently, the HUGO Gene Nomenclature Committee (HGNC) has a clear taxonomy for human mitochondrial gene names^10^. Still, anyone trying to analyze organelle genomes across different groups will be faced with a cornucopia of inconsistent names that can confound even experts in the field. In fact, recent large-scale comparative organelle genome analyses^6,7,11,12^ have repeatedly highlighted, in one way or another, the perils of and challenges brought by gene nomenclature disparities.

### Innovation

Here, we present the Gene Dictionary Tool (GDT), a protocol for the creation and implementation of a gene dictionary, which is comprehensive, versionable, and FAIR (Findability, Accessibility, Interoperability, and Reusability)^13^ -adherent. Unlike previous organelle gene dictionary attempts, GDT exhibits interoperability. Its main output, a .gdict file, has a parsable syntax and a structure that is inherently iterative. Detailed and human-readable log documentation lets any user make sense of the data. The standardized gene nomenclature in our .gdict files (available at our GitHub repository - https://github.com/brenodupin/gdt) is greatly reliant on the well-defined structure of the HGNC-approved mitochondrial gene symbols. The user can either adopt these suggested gene symbols (hereby referred to as gene labels) or change them as needed.

### Overview

The GDT protocol serves two main purposes: it creates a comprehensive gene dictionary from any genome(s) of interest while implementing a standardized gene nomenclature (that is chosen by the user). The gene dictionary creation process takes GFF3 files as input (with a .tsv file as their index) and, optionally, a .gdict file (be it “stripped” or “complete”). Creating the gene dictionary depends on retrieving identifying information from the respective GFF3 files and NCBI Gene queries, as well as input from the user (**Figure 1**). GDT does not correct annotations (it does not rely on BLAST); instead, it creates an expandable and customizable gene dictionary, allowing users to standardize gene names across disparate species. Note that we use the terms “gene” and “genome feature” interchangeably throughout this protocol (and in the GitHub page). That is because GDT can, in principle, analyze any annotated genome feature.

**Figure 1.**
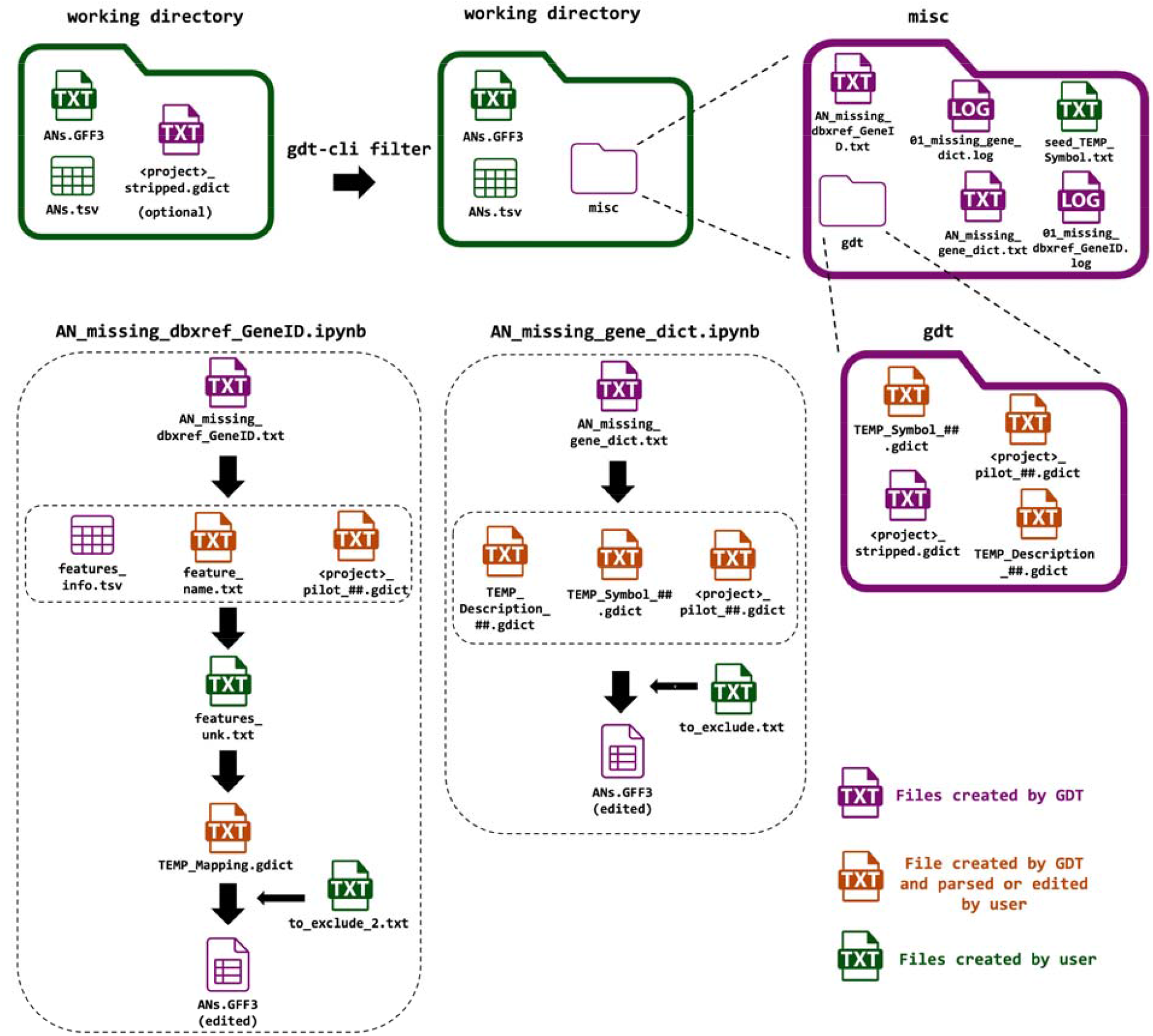
Overview of the GDT protocol. Creating and implementing a gene dictionary is a multipronged process relying on extensive user input. This diagram depicts all intermediate and final output files that can be generated by the protocol.

### Downloading the software and prerequisites

GDT is composed of four Python files, accompanied by two main Jupyter Notebooks. All files can be downloaded from GitHub (https://github.com/brenodupin/gdt). The Python files represent the gdt library, allowing users to interface with .gdict files (**Figure 2**). The accompanying Jupyter Notebooks (named “AN_missing_gene_dict.ipynb” and “AN_missing_dbxref_GeneID.ipynb”) let users iterate over NCBI Gene queries and create new .gdict versions. The library can be installed locally using pip (see the Step-by-Step methods for details), which will automatically install all necessary dependencies. For the Jupyter Notebooks, the user needs to install the necessary dependencies (e.g., biopython, pandas, and gdt) separately. The pipeline requires a Unix environment with Python 3.12 (or above). A third auxiliary Jupyter Notebook (named “Auxiliary_processes.ipynb”) enables users to update their .gdict files in a piecemeal fashion (see Troubleshooting).

**Figure 2.**
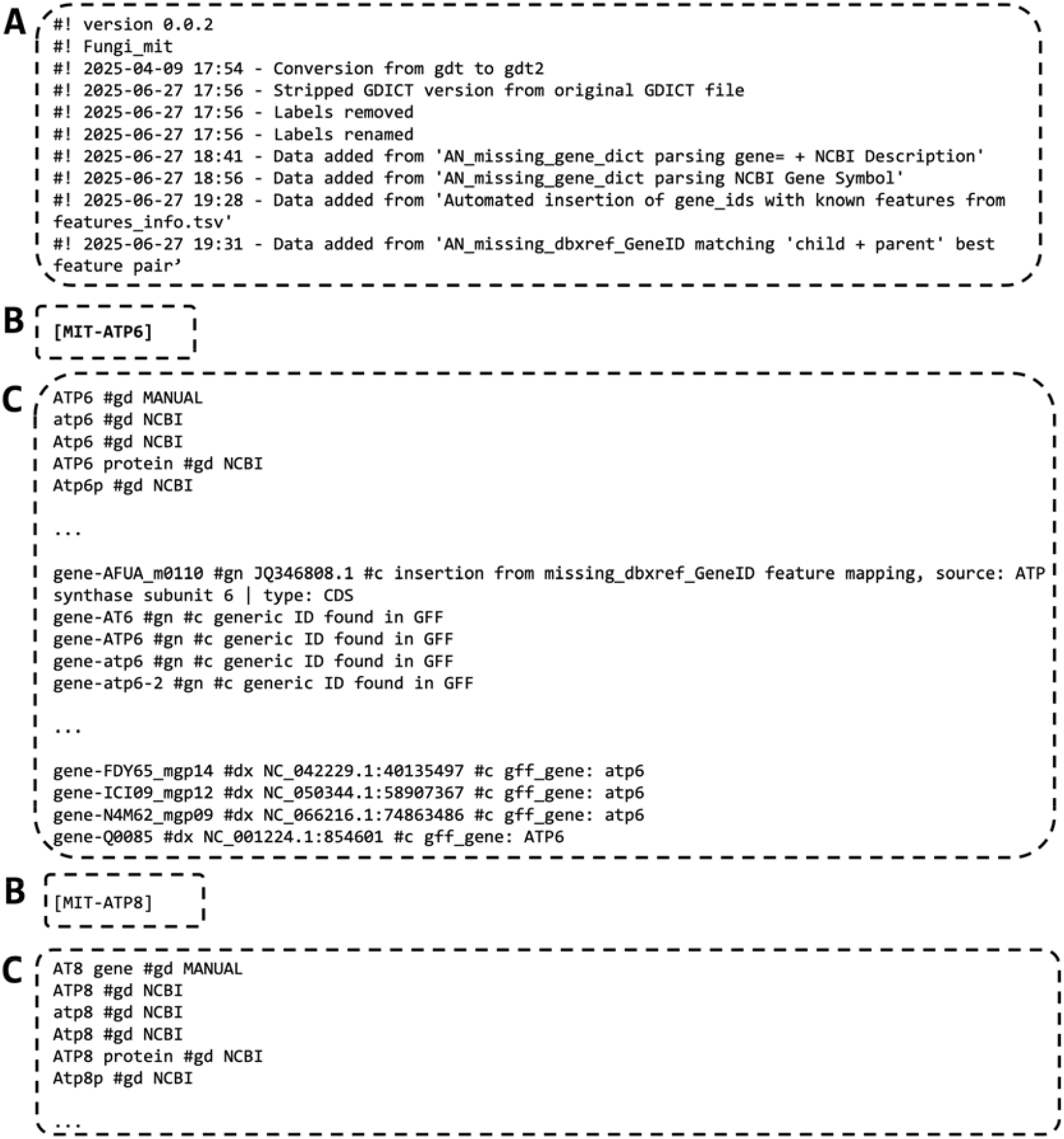
Example of a .gdict (Gene Dictionary) file (continues in the next page). (A) .gdict’s header section (B) .gdict’s label definition (C) .gdict’s data entries

***Note:*** specific hardware requirements, such as RAM memory and storage capacity, will depend on the size of the dataset. In principle, a common personal computer should have enough power to run GDT and analyze thousands of GFF3 files (of small genomes).

This example shows a .gdict file in its final version after multiple iterations through the protocol. A .gdict file is a plain-text file heavily inspired by the TOML structure. The .gdict structure is composed of three main elements: header section, label definition, and data entries. As a FAIR-informed file type, the header section contains metadata for traceability. Version number, time stamp, and the origin of the file are generated automatically by GDT. Group name (“Fungi_mit” in this example) is inputted by the user directly in the file. The actual dictionary starts right below the header and is read top-to-bottom, organized by gene labels (e.g., [MIT-ATP6]) and their respective data entries. For each gene label, genome features are classified (and sorted) according to the origin of their identifying information. The classification scheme is as such: “#gd” = GeneDescription, “#gn” = GeneGeneric, and “#dx” = DbxrefGeneID. The .gdict file also takes in commentaries in the “#c” field.

## Key resources table

| REAGENT or RESOURCE | SOURCE | IDENTIFIER |
| --- | --- | --- |
| Deposited data |  |  |
| Stripped and complete gene dictionary (.gdict) files | This manuscript | https://zenodo.org/records/1716776<br>6 |
| Examples of GFF3 files | This manuscript | https://zenodo.org/records/1716776<br>6 |
| Software and algorithms |  |  |
| gdt library, AN_missing_gene_dict.ipynb,<br>AN_missing_Dbxref_GeneID.ipynb,<br>Auxiliary_processes.ipynb. | This manuscript | [https://github.com](https://github.com/brenodupin/gdt)<br>/brenodupin/gdt |
| Python 3.12 | Python Software<br>Foundation | [https://www.pyth](https://www.python.org)<br>on.org |
| Biopython 1.85 | Cock et al.<sup>14</sup> | [https://biopython.](https://biopython.org/)<br>org/ |
| pandas 2.2.3 | The Pandas<br>Development Team | [https://pandas.py](https://pandas.pydata.org/)<br>data.org/ |
| Jupyter Notebook | Project Jupyter | https://jupyter.org<br>/ |
| NCBI-Gene | Sayers et al.<sup>15</sup> | https://www.ncbi.nlm.nih.gov/gene/ |

## Materials and equipment setup

The GDT tool is written in Python. The gdt library is composed of four Python scripts: cli.py, gdict.py, gff3_utils.py, and log_setup.py. The CLI (command-line interface) lets users interact with the gdt library in a user-friendly way (**Figure 3**). The two main accompanying Jupyter Notebooks (“AN_missing_gene_dict.ipynb” and “AN_missing_dbxref_GeneID.ipynb”) provide users with the functionalities necessary to build their own .gdict files iteratively. The only major external dependencies are pandas and Biopython. The NCBI Gene Database can be queried (extensively or not), depending on the type of information contained within the input dataset. All necessary dependencies are listed in the **KEY RESOURCES TABLE**. The source code for GDT is found at https://github.com/brenodupin/gdt.

**Figure 3.**
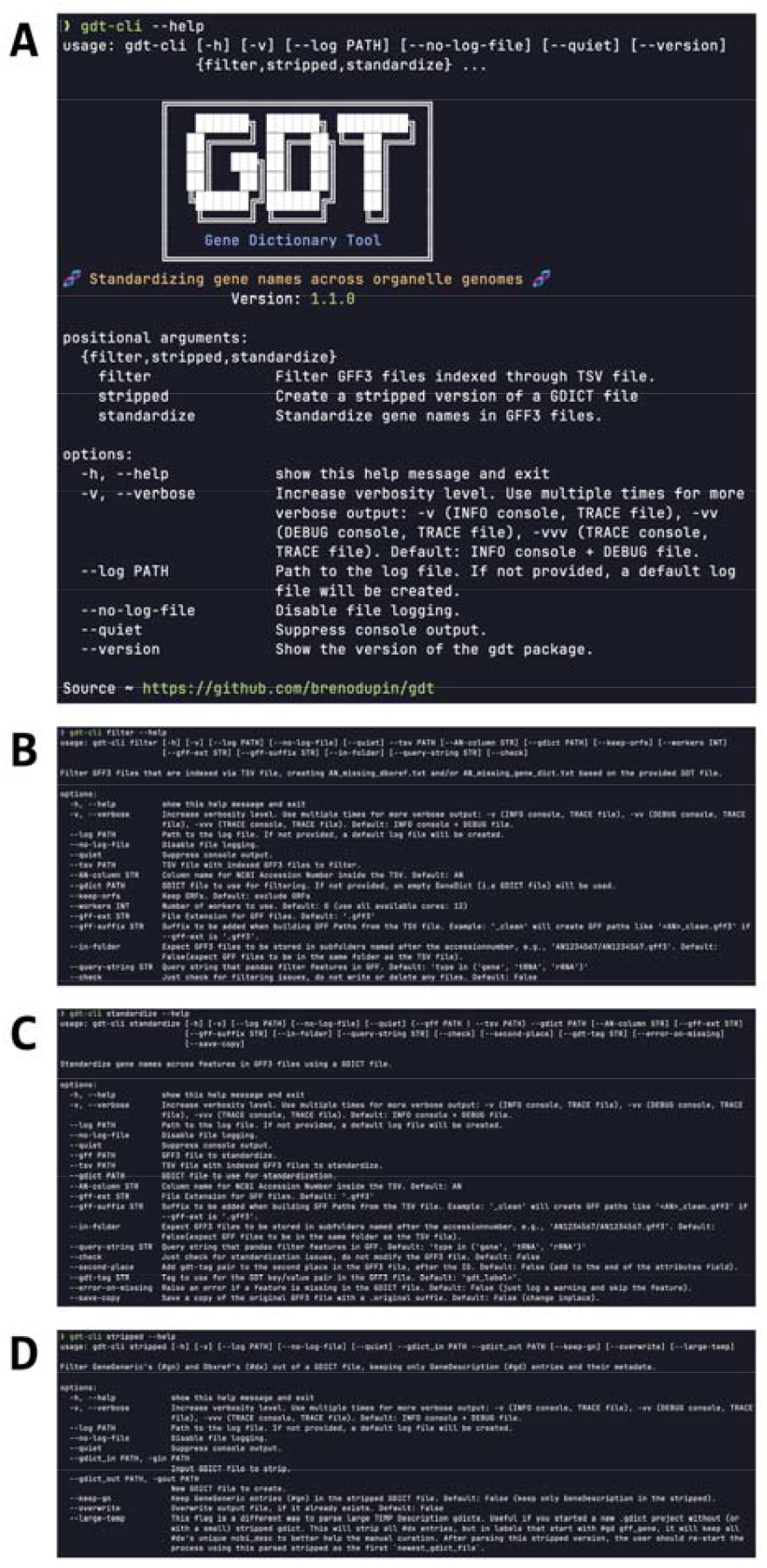
GDT’s command-line interface - CLI (continues in the next page). (A) GDT’s CLI help page. (B) gdt-cli filter –help output. (C) gdt-cli stripped –help output. (D) gdt-cli standardize –help output.

GDT’s CLI lets the user run the gdt library commands “filter”, “stripped”, and “standardize” without the need to edit the source code. This interface is extensively documented with detailed help messages to facilitate the use of GDT. The flag “–help” should be used anytime more information on GDT’s main functionalities is needed.

## Step-by-step method details

### Preparing the environment and files

This section prepares your local machine and files for the GDT protocol.

**Timing: ∼ 30 min**

1. Install GDT and Biopython using pip, as shown in the following command: `>pip install gdt biopython`
2. Download the Jupyter Notebooks from the “notebooks” folder in our GitHub page (https://github.com/brenodupin/gdt/tree/master/notebooks). The Notebooks should be saved in your chosen working directory.

***Note:*** you will need an IDE (Integrated Development Environment) to run the Jupyter Notebooks. For more information, consult Jupyter’s official documentation at https://docs.jupyter.org/en/latest/#where-do-i-start.

***Note:*** If you want to follow our example using the commands listed below, download fungi_mit.zip from the “example” folder of our GitHub (https://github.com/brenodupin/gdt/tree/master/example). In this case, unzip the folder with >unzip fungi_mit.zip and then enter the folder with >cd fungi_mit. From here, you can skip to step 4 to continue this protocol.

3. Compile a .tsv file with the NCBI accession numbers of your genomes of interest. Save this .tsv file in your chosen working directory and download all corresponding GFF3 files into this folder.

***CRITICAL:*** We have abbreviated “accession number” to AN in the .tsv file, but any other name can be used to refer to the accession number if the name is passed to the argument –AN-column in gdt-cli filter. The GFF3 files must be named as <AN.gff3>, where AN = “accession number” (as written in your .tsv file). The .gff3 extension can be changed (e.g., .gff) if the correct file extension is added to the flag –gff-suffix in GDT’s CLI, and Jupyter Notebooks. Consistency is key.

***Optional:*** Should you decide to build a dictionary from our stripped version (or from a previous version of any dictionary that follows the .gdict specifications), the .gdict file should also be added to the same working directory. The gdt-cli filter command will automatically move this .gdict file to the right subfolder for downstream analyses.

### Running gdt-cli filter –tsv –gdt

This section classifies the inputted genomes in three groups according to the presence of their genome features in the pre-existing dictionary: i) “ANs good to go”: all genome features of the given genome are already present in the gene dictionary; ii) “ANs missing gene_dict”: at least one genome feature is missing from the gene dictionary, and all genome features have dbxref IDs; iii) “ANs missing dbxref”: at least one genome feature is missing from the gene dictionary, and at least one feature does not have dbxref ID.

**Timing: ∼ 3 min (depending on dataset size)**

4. Run the gdt-cli filter command inside your working directory (**Figure 4**):

**Figure 4.**
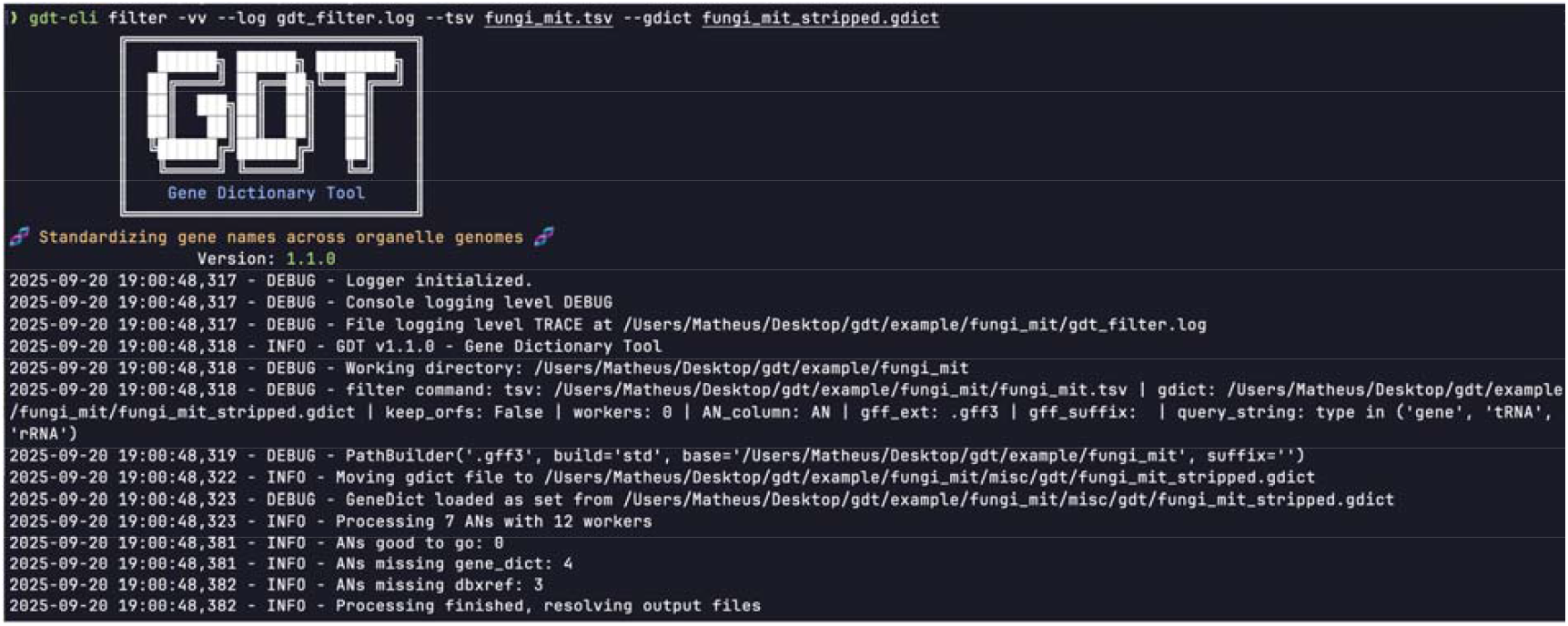
gdt-cli filter command example (continues in the next page).

>gdt-cli filter -vv --log gdt_filter.log --tsv fungi_mit.tsv --gdict fungi_mit_stripped.gdict

***Note:*** For a smoother experience, the command gdt-cli filter should be run in the terminal console from the working directory where the GFF3 and .tsv files are located. Alternatively, gdt-cli filter can be called from elsewhere, but –tsv and –gdt flags will take in either the absolute or relative paths to the respective files. In this case, the .tsv file’s parent folder will be considered the working directory. The gdt-cli has configurable verbosity levels (-v, -vv, and -vvv) for debugging.

***Note:*** *For each accession number listed in the .tsv file, gdt-cli filter will scan through the respective GFF3 file and try to match its genome features to those listed in the inputted .gdict file. Assuming that you started your analysis with an incomplete gene dictionary (e.g., a stripped .gdict), some of your GFF3 files will likely have genome features that are not listed in your initial .gdict file. There are then two possibilities: a given accession number will be added to AN_missing_gene_dict.txt only when all genome features of the GFF3 file have Dbxref GeneIDs. If at least one genome feature does not have Dbxref GeneID, the corresponding accession number will be listed in AN_missing_dbxref_GeneID.txt. These two files will be used subsequently in the accompanying Jupyter Notebooks (each named after the gdt-cli filter output file names)*.

***CRITICAL:*** The default genome feature types that will be searched in each GFF3 file are “gene”, “tRNA”, and “rRNA”. Should you want to change the default (e.g., to search “gene” only), you can change the –query-string flag as in: “type == ‘gene’” or “type in (‘gene’, ‘tRNA’)”.

In this example, the verbosity level -vv is used. Note the detailed (and user-friendly) logging that is provided when -vv is active. The -vv level will print onto the console screen log information at DEBUG level and save log information into a separate log file at TRACE level. Immediately after running gdt-cli filter, the user will know i) the number of genomes (i.e., ANs = accession numbers) for which all chosen features are already listed in the inputted .gdict, ii) how many have genome features that are not listed in the gene dictionary (but have Dbxref GeneIDs; “ANs missing gene_dict:”), and iii) how many have features that are not in the gene dictionary and lack Dbxref GeneIDs (“ANs missing dbxref:”).

### Running AN_missing_gene_dict.ipynb

This section queries through NCBI Gene (https://www.ncbi.nlm.nih.gov/gene/) to add the inputted genome features into the (most likely) appropriate gene symbols.

**Timing: ∼ hours to days (depending on dataset size and time spent in manual curation)**

5. Open the first Jupyter Notebook (AN_missing_gene_dict.ipynb) in your IDE (Integrated Development Environment) of choice – e.g., VS Code.
6. Run the first cell (Imports and functions) to import all necessary dependencies and instantiate all required global variables and related functions. ***Note:*** In both Jupyter Notebooks, we recommend running each cell sequentially (i.e., one by one) instead of all at once. This will ensure that processing errors are caught and will help prevent the program from crashing.
7. Edit the following variables in cell A. of the Setup section: DATA_DIR, newest_gdict_file, global_query_string, remove_orfs, gct, gff_sufix, Entrez.email, and Entrez.api_key. ***Note:*** global_query_string is pre-defined in the gdt library with two possible values: QS_GENE (“type == ‘gene’”) or QS_GENE_TRNA_RRNA (“type in (‘gene’, ‘tRNA’, ‘rRNA’)”). This variable will determine what type(s) of genome features will be analyzed. To add a new genome feature (e.g., exon or ncRNA), insert the new type name to the list that defines this variable.

***CRITICAL:*** if you change global_query_string, you must restart the process. The new global_query_string should be passed to gdt-cli filter as the flag –query-string.

a. Update newest_gdict_file, as you iterate through the protocol and update your working .gdict. This variable represents the latest iteration of the gdt creation process, and it is highly recommended that you start your analysis with the _stripped.gdict version (of the appropriate organism group) provided in the GitHub page.

***CRITICAL:*** When using the stripped version of GDT, the file name must contain the suffix “stripped” (as in fungi_mit_stripped.gdict), otherwise the protocol will crash. The “newest_gdict_file =” variable only contains the name of the respective file inside the gdt folder. This is not mandatory, except in the first pass of either AN_missing_gene_dictionary.ipynb or AN_missing_dbxref_GeneID.ipynb. Whenever a new version of the .gdict file is being created, you must update “most_recent_gdict_filename =” to the latest iteration of your .gdict and run cell A. of the Setup section again. This applies to both Jupyter Notebooks.

***Note:*** gct stands for “genetic compartment”. In its current version, we use “MIT” for mitochondria and “PLT” for plastid. These acronyms are added as prefixes to the gene labels, e.g., [MIT-ATP6]. You can change this prefix as you see fit. Entrez.email and Entrez.api_key are private and specific to each user. These values are obtained from the user’s NCBI account. The email is mandatory, regardless of the size of the dataset, whereas the API key is highly recommended when analyzing large(r) datasets because it gives the user a higher API request rate.

8. Prepare a (mandatory) logger for debugging in cell B. of the Setup section. Log files are stored in the misc folder. If you do not want a log file, change the “print_to_console=” and “save_to_file=” to False.
9. Run all cells (from A. to E.) in the section “TEMP First Pass”. ***Note:*** Cell A.: the AN_missing_dict.txt will be loaded. Cell B.: the .gdict file (stored in the most_recent_gdict_filename) will be loaded onto the variable gene_dict. Cell C: the next cell composes the core process of this step. Briefly, each GFF3 file will be loaded and scanned for the chosen features (e.g., “gene”). If the corresponding feature ID (found in “ID=“) is already present in the loaded .gdict, the feature is skipped, otherwise the values of “gene=” (or “Dbxref=GeneID”) will be used to match the respective feature to .gdict’s gene labels.

***Pause point:*** after having run cell C., you should pause and check whether any errors have been listed at the end of your log file. These errors can be of type “File not found” or “Entrez.read”. The log messages will help you troubleshoot.

***Note:*** Cells D. and E. write and save the temporary gene dictionary (temp_gene_dict) and the latest iteration of the gene dictionary (gene_dict) (**Figure 5**).

**Figure 5.**
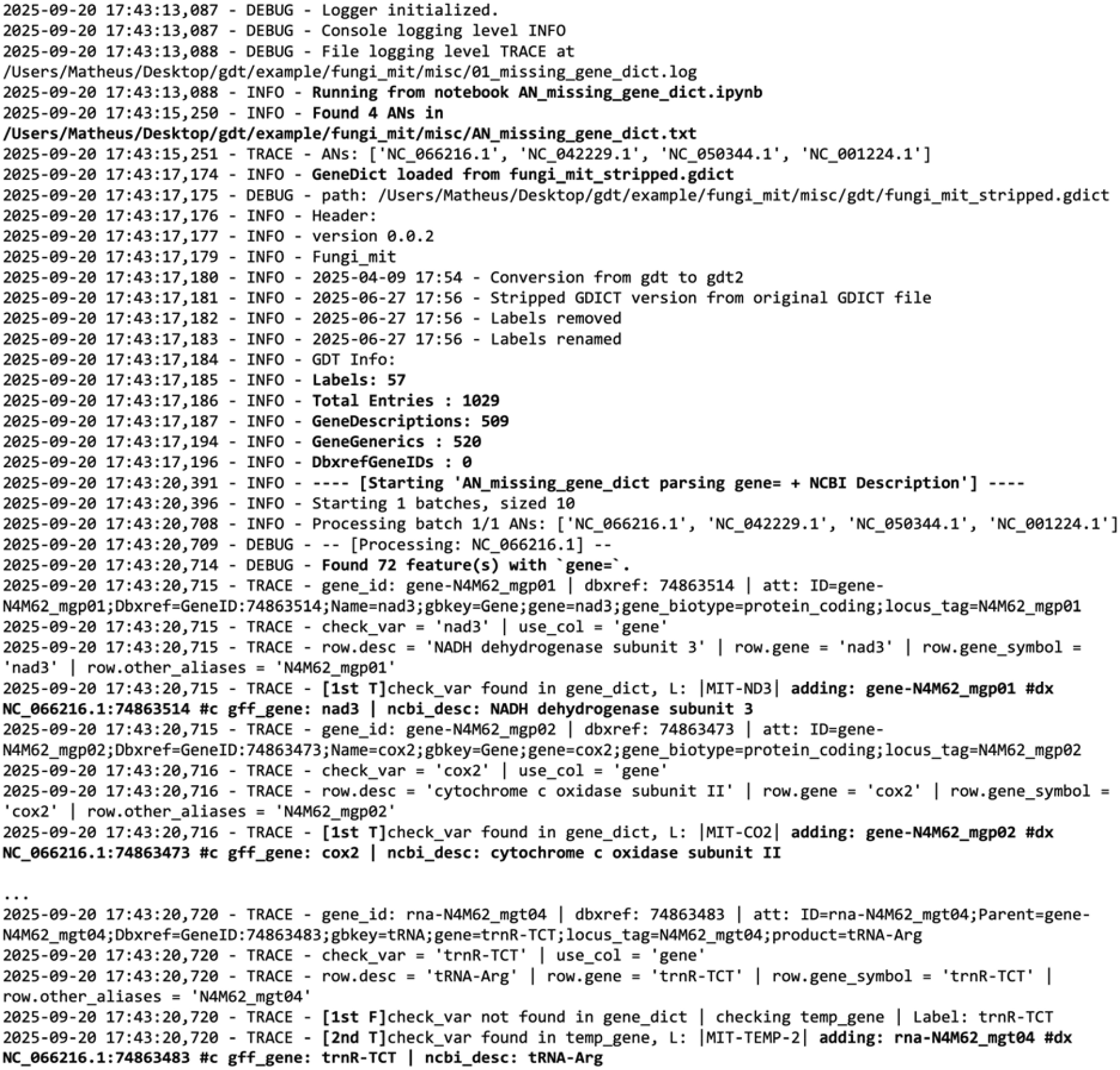
initial output log file of AN_missing_gene_dict.ipynb (continues in the next page).

Example of a human-readable and machine-parsable log output showing the user how each genome feature was parsed. Note the detailed header describing what .gdict iteration is being loaded (stripped or pilot_XX), how many genomes are being analyzed (in this case, 4 accession numbers listed in AN_missing_gene_dict.txt), and the structure of the loaded dictionary, including the number of entries (gene dictionary length), number of standardized gene labels, and number of #gd, #gn, and #dx entries. The genome features that are left unmatched will be added to the TEMP_Description_XX.gdict file (see next step).

10. Parse manually TEMP_Description_##.gdict (**Figure 6**).

**Figure 6.**
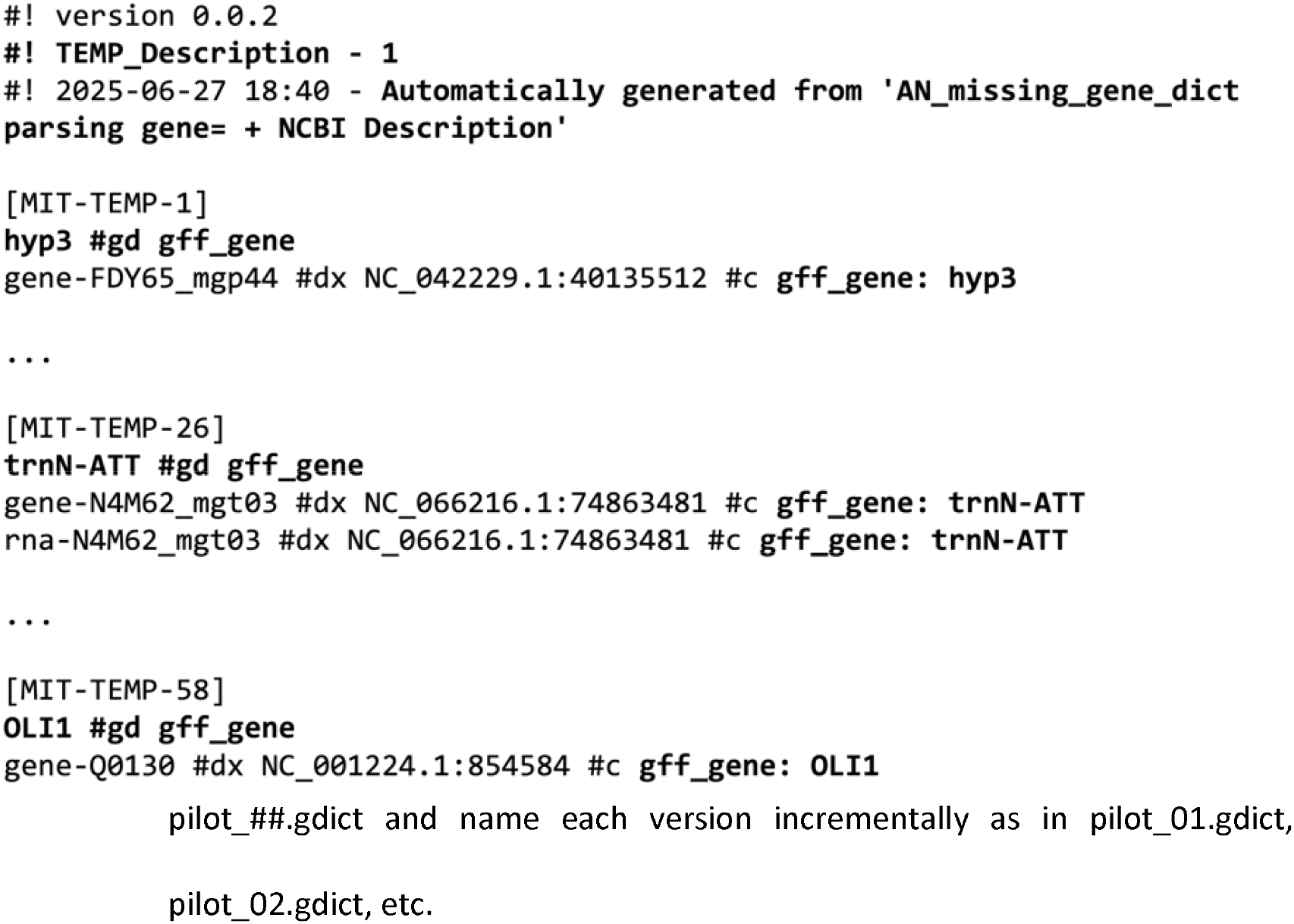
Excerpts of our TEMP_Description_01.gdict and its respective log information. Just like any other .gdict file, the TEMP_Description_XX.gdict has a time-stamped incremental header, which indicates when and how this file was created (i.e., automatically from AN_missing_gene_dict.txt).

***CRITICAL:*** If you can classify the genome feature with the information obtained from “gff_gene” and/or “ncbi_desc”, you should copy the respective line(s) from TEMP_Description_##.gdict to your latest .gdict. If you cannot classify the genome feature with the provided information, you should copy the line(s) from TEMP_Description_##.gdict to a new file named seed_TEMP_Symbol.txt.

***Note:*** For better traceability, we suggest that you maintain the output files as they are created and create/save new files as you progress through the analysis. As you iterate through the protocol and manually add new entries to your .gdict files, we highly recommend that you to save each new pilot_##.gdict and name each version incrementally as in pilot_01.gdict, pilot_02.gdict, etc.

11. Update the most_recent_gdict_filename to the latest .gdict you just edited. Then, run the “Setup” (cell A.) again.
12. Run “TEMP Second Pass” section, if you end up having the seed_TEMP_Symbol.txt. ***Note:*** This process follows the same rationale as described in item 8. The only difference is that the identifying information is obtained from “NCBI gene symbol” field, instead of “NCBI gene description” field. The two final cells of this section will write (and save) a temporary gene dictionary (temp_symbol_gene_dict) and automatically update the latest gene dictionary (gene_dict).
13. Parse manually TEMP_Symbol_##.gdict. ***Note:*** The rational here is the same as in item 9. However, unclassifiable genome features should either be added to a new file called to_exclude.txt or the whole genome should be removed from the dataset.
14. Run cells A. and B. of the final section (“Gene exclusion of to_remove.txt), if you end up creating to_exclude.txt. ***Note:*** This step adds (directly into the original GFF3 files) the tag “discard-” before the genome feature(s) to be excluded. This guarantees that these features will not be included in any future iteration of GDT.

### Running AN_missing_dbxref_GeneID.ipynb

This section parses the genome features that do not have a dbxrefID value by extracting identifying information from the corresponding GFF3 files. This information is used to bin the genome features according to the gene labels in the dictionary.

**Timing: ∼ hours to days (depending on dataset size and time spent in manual curation)**

15. Run AN_missing_dbxref_GeneID.ipynb, if gdt-cli filter created AN_missing_dbxref_GeneID.txt.
16. Define other essential variables and the logger, in the Setup section (just as you did at the beginning of AN_missing_gene_dict.ipynb). ***Note:*** See items 6 and 7 of “Running AN_missing_gene_dict.ipynb” for a refresher on what each variable means.
17. Run cells A to D of the “Features Extraction” section. ***Note:*** Briefly, the AN_missing_dbxref_GeneID.txt is read, the latest .gdict file is loaded, and the respective GFF3 files (of the accession numbers listed in AN_missing_dbxref_GeneID.txt) are analyzed. The attributes column of each genome feature will be broken down according to a series of regular expressions that are defined at the beginning of this script. The output will be “features_info.tsv” and “feature_names.txt”. The last column of “features_info.tsv” (named “best_feature”) will contain the attribute information deemed to be most relevant for subsequent parsing. This column will give rise to “feature_names.txt”. “best_feature” derives from the variable “features_order”, which is defined in “Imports and functions” section of this script. The features_order is a list of attributes ranked according to their discriminatory power. Based on our experience with organelle genomes, we have chosen the following order to be the default: ‘features_order = [“gene”, “product”, “description”, “name”, “note”, “gene_synonym”]’. You have the flexibility to change the order and/or add new attributes to this variable.

***Pause point:*** After having run cell C. in “Features Extraction”, pause and check whether an error message appears. Detailed log messages will guide you through the troubleshooting.

***Note:*** In cell D., there exists a simple “tRNA parser” to preprocess tRNA annotations. This parser is based on the patterns of the usual tRNA gene nomenclature scheme in organelle genomes. The goal of the parser is to automate (as much as possible) to subsequent parsing of genome features. See Limitations for more information.

18. Parse manually “feature_names.txt”.

***CRITICAL:*** Check whether any entries can be added to your latest .gdict. Once you add these entries to .gdict, save the new .gdict version. If you have entries that are still unidentifiable, you should create a new txt file in the misc folder (named feature_unks.txt) and add the unidentifiable entries to it. After having created and saved your new .gdict, update newest_gdict_file in the Setup section and rerun the respective cell.

19. Run cell A in the “Automated Insertion from features_info.txt” section. ***Note:*** This cell will check whether any entries from features_info.tsv, that are not in feature_unks.txt, are missing in the latest .gdict. If that is the case, you will receive a warning (in the log and/or screen) for what features should be double checked. The warning message alert you of three possibilities: 1) you forgot to add the respective entries to your latest .gdict, 2) you forgot to reload the latest .gdict, or 3) you forgot to add the entries to “feature_unks.txt”. After addressing the issues identified in this step, run this cell again to ensure that that there are no remaining issues.
20. Run cells B. and C. of this section. ***Note:*** Cell B.: this step will scan features_info.tsv and add to your latest .gdict all genome feature IDs that have a “best feature” that is not listed in “feature_unks.txt”. Genome feature IDs will be added under the corresponding gene labels according to their “best_feature”. Cell C.: a new .gdict iteration is saved with an updated header containing the respective traceable information (i.e., a stime stamp, iteration number, etc).

***CRITICAL:*** you *must* ensure that your newest_gdict_file is updated in the Setup section and then rerun this section before moving to the next step.

21. Run the section “TEMP Mapping” from A. to D. ***Note:*** Cell A: will reload the latest .gdict. and will extract the species and their corresponding feature IDs that need to be more investigated. This extraction happens through the filtering of “best_feature” that are found in “feature_unks.txt”. Cell B: as a final attempt to find descriptive information about these features, this step will (try to) map child features to their corresponding parent features. Through this mapping, the descriptive information of the child feature will be relayed to the parent feature. Once mapped, the parent feature is compared to the latest .gdict and, if there is a match, added to it. If there is no match, the parent feature will be added to TEMP_Mapping for manual parsing. The features that could not be mapped to any child are also added to TEMP_Mapping but are marked with the - UNKNOWN label. Cells C. and D: TEMP_Mapping and the new iteration of .gdict will be saved, if applicable.
22. Parse TEMP_Mapping and add any further entries to your .gdict. Add entries that remain unidentifiable to “to_remove_2.txt”. ***Note:*** This parsing is similar to the process described in items 9 and 12, but “to_remove_2.txt” should not be confused with “to_remove.txt” (which was created at the end of AN_missing_gene_dict.ipynb). Both files serve the same purpose, but for different steps of the protocol. Just as with “to_remove.txt”, you should create “to_remove_2.txt” as an empty file in the misc folder.
23. Run cells A. and B. in the final section (“Gene exclusion of to_remove_2.txt), if you end up creating to_exclude_2.txt. ***Note:*** this step is similar to step 13.

***Optionally:*** To verify if there are any unaccounted gene IDs left in your dataset, run gdt-cli filter one last time using your most complete (i.e., latest iteration of) .gdict. If there are any unidentified genome features, gdt-cli filter will create the appropriate file(s) (either AN_missing_gene_dict.txt or AN_missing_dbxref_GeneID.txt) and the process cycles over again.

***Optionally:*** if you want to add the standardized gene labels to your GFF3 files, run gdt-cli standardize.

## Expected outcomes

The main outcome of this protocol is a comprehensive and versionable gene dictionary saved as a .gdict file. The dictionary is not only intuitive and human readable but easily parsable by a computational agent. Along with the .gdict file, this protocol generates intermediate files (both of type .gdict and .log) that provide a detailed view of the analysis and permit users to act on the data as needed. The protocol also gives the option to add standardized gene dictionary labels to the inputted GFF3 files for downstream analyses. Users can interact with our protocol in two main ways: i) they can create their own gene dictionaries from their genomes of interest (with the possibility of creating their dictionary gene labels); or ii) they can apply the gene dictionaries created by us and implement the protocol as a way to bring consistency to the gene nomenclature across (organelle) genomes of their choosing. Another contribution of this protocol (regardless of whether one creates their own dictionary or deploys ours) is the launching of the gdt library. This Python library, with its user-centered design, enables users to manipulate and integrate .gdict files into other pipelines seamlessly. We hope that these outcomes will contribute to a more consistent and FAIR-guided gene nomenclature ecosystem.

## Limitations

Most (if not all) of the limitations we have identified in our analyses are applicable to any type of annotated genome. The first limitation pertains to the time spent in manual curation. Although the dataset (i.e., the number of GFF3 files to be analyzed) might increase linearly, the final number of genome features to be parsed is poised to grow much faster. This is because each genome will likely add several genome features to the dataset. This limitation is ameliorated as the user iterates through the pipeline and creates a more complete version of their working .gdict. To provide users with a head start on their analysis of organelle genomes, the stripped and complete version of our .gdict files are available in our GitHub page (https://github.com/brenodupin/gdt/tree/master/resources).

Another limitation comes from the inherent structure of GFF3 files. The Generic Feature Format (GFF) file has become a staple in genome analysis because major stakeholders like NCBI, and Ensembl release their genome annotations in the GFF3 format (notwithstanding other formats like BED). The difficulties (and frustration) that come with trying to analyze GFF3 files from disparate sources have been shared in popular media outlets like Medium (https://medium.com/@shiansu/the-not-so-standard-gff3-standards-957c54d848af) and in The-Sequence-Ontology GitHub Specifications page (https://github.com/the-sequence-ontology/specifications/blob/master/gff3.md). Upon reading the current specifications (version 1.26, released on 18 August 2020), one can clearly find two potential pitfalls: the first is that IDs are mandatory only for features that possess children features (such as protein-coding genes). As GDT’s framework relies on unique genome feature IDs, annotations without IDs cannot be indexed in the .gdict file and are rendered incompatible with our protocol. The second pitfall is that “IDs for each feature must be unique within the scope of the GFF file”, so these IDs are guaranteed to be unique only within each GFF3 file. In other words, a single ID can refer to a different genome features in different genomes (e.g., the same ID can refer to ATP9 in one genome and to ATP8 in another). We expect this potential pitfall to be more likely to occur when the user analyses different annotation versions of the same genome (as in NC_??######.1 and NC_??######.2, where .1 and .2 represent different versions of the same genome) or when a single genome has different chromosomes. This issue also becomes more probable in very large datasets. In fact, in our analyses of thousands of organelle genomes, we have found two cases of “genome feature ID collision” – one in chloroplast and one in mitochondrial genomes. In this situation, we removed the genomes from our analysis. See the Troubleshooting section for how the protocol handles a somewhat similar collision issue and how one can update the .gdict to add new versions of a given genome annotation.

Having only tested our protocol on organelle genomes can be seen either as an asset or as a limitation. Organelle genomes are notorious for their unconventional genomic attributes and inconsistent annotations. Arguably, we might have picked the toughest case scenario, which has forced us to adopt defensive programming from top to bottom. But we can only script defensively against what we have encountered (and/or predicted), so testing GDT on other types of genomes is an obvious next step in the development of our protocol. In its current form, GDT relies only on NCBI Gene queries when searching (externally) for further information on the annotation of the genome feature at hand. We do that through the unique NCBI gene IDs that are assigned to “Dbxref=” in the GFF3 attributes column. Another further improvement of this protocol is to give the user the possibility to tweak the cross-reference search to a database of their choosing.

After having run this protocol on thousands of organelle genomes, some clear trends within the “hard-to-identify” genome features have surfaced. Features annotated as “hypothetical protein” and “orf” are discussed in the Troubleshooting section, but tRNAs deserve their own mentioning within limitations. Given the myriad of tRNA gene naming styles, one should expect to have to manually parse tRNA names. For example, issues vary from “trnA vs tRNA” (the first referring to tRNA-Ala and the second referring to a generic tRNA) to “gene-trnF(GAA) vs gene-trnF (GAA)” and anything in between. To a certain extent, consistency can be distilled from organelle tRNA names, so much so that the use of regular expressions is commonplace when dealing with tRNAs. But variability and discordance within organelle tRNA nomenclature have been extensively highlighted^8^, even though clear guidelines have been proposed over four decades ago^9^. We have added a basic tRNA-parser capability to GDT’s latest version (in AN_missing_dbxref.ipynb). This basic parser has been designed according to the patterns we have identified in our analyses, so we invite future users to try and extend the parser according to their needs.

## Troubleshooting

### Problem 1

What if I do not want to adopt the gene labels proposed in the GDT protocol?

### Potential solution

You have full autonomy to create your own gene labels. This protocol lets users create a comprehensive gene dictionary regardless of how the dictionary is labeled.

### Problem 2

I have hundreds of “hypothetical protein” features in my TEMP_Description_##.gdict. What should I do? Similarly, what should I do with very generic/cryptic entries like “rna”, “trna”, “hyp”, or “oi”?

### Potential solution

We have two suggestions: you can either create an “umbrella label” for all these problematic genome features (such as [MIT-HYP] for mitochondrial hypothetical proteins) or you can divert them directly to “TEMP Symbol processing” to obtain more descriptive information. In our own experience, we have found some interesting genome features buried in the “hypothetical protein” pile. Sending these features to Temp Symbol will likely clutter your TEMP_Symbol.gdict, so feel free to use to_remove.txt option as well.

### Problem 3

What do [1st T], [1st F], [2nd T], and [2nd F] mean in the log file?

### Potential solution

[1st T], [1st F], [2nd T], and [2nd F] serve as signposts in the staggered parsing that happens when the data_process function is called. The question being asked is always “is this genome feature already in my .gdict”?, but data_process asks this question (three times) in slightly different ways depending on where you are in the analysis.

[1st T]: genome feature ID’s check_var is in the current gene dictionary (i.e., gene_dict), where check_var can be the information from gene, NCBI gene symbol, or NCBI gene description (the source of check_var is defined in use_col). Then, the genome feature ID is added to the current gene dictionary immediately under the check_var’s gene label.

[1st F]: check_var is not in gene_dict (so go to the question below).

[2nd T]: check_var is in TEMP_Description_##.gdict (or TEMP_Symbol_##.gdict).

[2nd F]: check_var is not in TEMP_Description_##.gdict (or TEMP_Symbol_##.gdict).

### Problem 4

After having run cell B. in “Automated Insertion from features_info.txt”, I got the error ‘Multiple best_features with different labels’. What should I do?

### Potential solution

This error occurs when the script identifies a gene_id that has multiple ‘best_features’, and these features, when queried in the .gdict, points to different labels. Although the use of the same feature ID in different GFF3 files is somewhat common (e.g., “ID=gene-atp9;”) for files without Dbxref GeneID, this ID collision becomes a problem when the IDs refer to incongruent features {e.g., “ID=rps;”). The solution is to manually edit one of the GFFs to differentiate one of the gene_IDs, then edit the features_info.tsv to reflect the change (or just run the entire AN_missing_dbxref_GeneID.ipynb again). It is also possible to just remove the offending genome from your dataset.

### Problem 5

One of my GFF3 files got a new version. What should I do?

### Potential solution

When a new version of a GFF3 file is released, and you would like to update your gene dictionary accordingly, you can run the “Removal of an GFF’s file annotations from a GeneDict” using Auxiliary_processes.ipynb. Detailed guidelines on how this functionality works can be found in our GitHub (https://github.com/brenodupin/gdt/tree/master/notebooks).

## Resource availability

### Lead contact

Further information and requests for resources should be directed to and will be fulfilled by the lead contact, Matheus Sanita Lima.

### Technical contact

Technical questions on executing this protocol should be directed to and will be answered by the technical contact, Breno Dupin.

### Material availability

not applicable.

### Data and code availability

The datasets and code generated during this study are available at GitHub: https://github.com/brenodupin/gdt and Zenodo: https://zenodo.org/records/17167766.

Acknowledgments

The authors wish to acknowledge the Natural Sciences and Engineering Research Council of Canada (NSERC) and the Fundação Araucária for funding granted to D.R.S. (Discovery Grant, NSERC) and to A.R.P. (NAPI Bioinformática 66.2021, Fundação Araucária).

## Author contributions

Conceptualization: M.S.L., B.D., D.R.S., and A.R.P. Software and Methodology: B.D. Data curation: M.S.L. and B.D. Writing – original draft: M.S.L. and B.D. Writing – review & editing: M.S.L., B.D., D.R.S. and A.R.P. Funding acquisition and Supervision: D.R.S. and A.R.P.

## Declaration of generative AI and AI-assisted technologies in the manuscript preparation process

During the preparation of this work the author(s) used Claude AI and ChatGPT for script optimization and documentation, code formatting, GitHub backend configuration, tool logo generation and alike. After using this tool/service, the author(s) reviewed and edited the content as needed and take(s) full responsibility for the content of the published article.

